# Interleukin-11 plays a key role in human and mouse alcohol-related liver disease

**DOI:** 10.1101/2021.08.24.456749

**Authors:** Maria Effenberger, Felix Grabherr, Benedikt Schaefer, Christoph Grander, Lisa Mayr, Julian Schwärzler, Barbara Enrich, Patrizia Moser, Julia Fink, Alisa Pedrini, Nikolai Jaschke, Martin Freund, Alexander Loizides, Reto Bale, Daniel Putzer, Anissa A Widjaja, Sebastian Schafer, Stuart A Cook, Heinz Zoller, Georg Oberhuber, Timon E Adolph, Herbert Tilg

## Abstract

**Background:** Alcoholic hepatitis (AH) reflects acute exacerbation of alcoholic liver disease (ALD) and is a growing healthcare burden worldwide with limited treatment options. Interleukin-11 (IL-11) is a pro-fibrotic, pro-inflammatory cytokine with increasingly recognized toxicities in parenchymal and epithelial cells.

**Aim:** The aim of this study was to explore the prognostic value of IL-11 serum levels in patients suffering from AH and cirrhosis of various etiology and to understand the role of IL-11 in experimental ALD.

**Methods:** IL-11 serum concentration and tissue expression was determined in a cohort comprising 50 patients with AH, 110 patients with cirrhosis and 19 healthy volunteers. Findings were replicated in an independent patient cohort including 186 patients. Ethanol-fed wildtype mice were treated with a neutralizing murine IL-11 receptor-antibody (anit-IL11RA) and thereafter examined for severity signs and markers of ALD.

**Results:** Human IL-11 serum concentration and liver tissue expression increased with severity of liver disease and were most pronounced in AH. In a multivariate Cox-regression, a serum level above 6.4 picograms/milliliter was a MELD independent risk factor for transplant-free liver disease survival in patients with compensated and decompensated cirrhosis. Findings were confirmed in an independent cohort. In mice, severity of alcohol-induced liver inflammation was positively correlated to enhanced hepatic IL-11 expression. Pretreatment with a neutralizing anti-IL11RA inhibited hepatic inflammation and mice were protected from ethanol-induced liver injury. In comparison to IgG-control, ethanol-fed mice treated with anti-IL11RA showed decreased steatosis, hepatic neutrophil infiltration, and expression of pro-inflammatory cytokines.

**Conclusion:** IL-11 plays a crucial role in the pathogenesis of ALD and could serve as an independent prognostic factor for transplant-free survival. Blocking IL-11 signaling might be a therapeutic option in human ALD, particularly AH.

## Introduction

Chronic alcohol consumption causes physical and psychiatric disabilities and contributes to approximately 3 million deaths each year worldwide. Overall, ethanol abuse is responsible for 5.1% of the global burden of disease^1^. Alcoholic liver disease (ALD) is a common liver disease and is a complex pro-inflammatory process leading to steatosis, alcoholic hepatitis (AH), fibrosis, cirrhosis, and finally hepatocellular carcinoma^2^. The pathogenesis of ALD, especially AH, is poorly understood which is reflected by poor therapeutic options and patient outcome. Along with direct toxicity of ethanol and acetaldehyde, alcohol causes gut dysbiosis, which subsequently provokes an altered gut barrier function and endotoxemia^3-5^. This dysregulation also induces expression of various pro-inflammatory cytokines in the liver which contribute to cell death and liver injury^6^. Interleukin (IL)-6, tumor necrosis factor-α (TNF-α), and IL-1β are all important for ALD in experimental models^7-9^.

IL-11 is one of eleven known IL-6 family members^10,11^. It is secreted from epithelial cells and hepatocytes in response to injury and from activated fibroblasts, acting in an autocrine and paracrine manner. In epithelial cells and hepatocytes, IL-11 causes cellular dysfunction and can initiate cell death processes while blocking regeneration^12,13^. In stromal cells, IL-11 triggers extracellular matrix production and invasion and migration of myofibroblasts^14^. To complete this vicious cycle, activated fibroblasts and myofibroblasts then secrete pro-inflammatory cytokines and chemokines^15^. All members of the IL-6 family require the ubiquitously expressed gp130 receptor to activate downstream signaling pathways. IL-11 was often compared to IL-6, as both cytokines form a similarly arranged gp130 heterodimer complex to initiate cell activation. Interestingly, the α-subunit receptors of these two cytokines are expressed in almost mutually exclusive cell types and IL-11 and IL-6 are proving to have very different functions^16^.

Little is known about the role of IL-11 in liver disease and its impact on inflammation of the liver. Recently published experiments showed a central role for endogenous and species-matched IL-11 in the pathology of fibro-inflammatory diseases, such as inflammatory bowel disease, myocardial and lung fibrosis, systemic sclerosis, and, most importantly, non-alcoholic liver disease where direct hepatotoxic effects have been shown^16-19^.

We investigated circulating IL-11 and its expression in the liver in a patient cohort suffering from cirrhosis and AH and establish the concept of IL-11 signaling as driver of experimental ALD.

## Materials and Methods

### Patients

Patient’s samples of the training cohort (n=179) were collected at the University Hospital of Innsbruck between 2017 and 2019, whereas samples of the validation cohort (n=186) were collected between 2019 and 2020.

Cirrhosis was confirmed by abdominal computed tomography and by indirect cirrhosis signs, such as esophageal varices, portal hypertension, ascites, hepatic encephalopathy, or thrombocytopenia. The Model of End stage Liver Disease (MELD) score was calculated. Distribution of etiology of liver disease in the training group (n=160) was as follows: 79 patients had compensated cirrhosis and 31 patients had decompensated cirrhosis. The underlying etiology of liver disease is summarized in Table 1. Decompensated cirrhosis was defined by presence of ascites, bleeding, encephalopathy, or jaundice^20^. Patients diagnosed with cirrhosis due to alcoholic fatty liver disease and confirmed alcohol abstinence > 6 months were included in the cirrhosis group. Another group in the training cohort consisted of patients with alcoholic hepatitis (AH) (n=50) which was diagnosed upon an acute onset of jaundice in the past 8 weeks arising on the background of severe alcohol abuse. The thresholds was defined at an average alcohol consumption of more than 3 drinks (40 grams (g))/day for women and 4 drinks (50-60 g)/day for men. Patients diagnosed with AH were addicted for at least 6 months with no abstinence until 60 days before the onset of jaundice. The diagnosis of AH required a serum bilirubin > 3 mg/dl and serum aspartate transaminase (AST) > 50 IU/ml. It’s ratio to alanine aminotransferase (ALT) had to be above 1.5. Patients with AST and ALT levels > 400 U/ml were excluded from the alcoholic hepatitis group^21^.

**Table 1:** Patient’s characteristics of the training cohort.

|  | **Healthy**<br>**(n=19)** | **compensated**<br>**cirrhosis**<br>**(n=79)** | **decompensated**<br>**cirrhosis (n=31)** | **alcoholic**<br>**hepatitis**<br>**(n=50)** |
| --- | --- | --- | --- | --- |
| **Age (years)** | 61 ($\pm 13.1$) | 63.5 ($\pm 9.7$) | 57 ($1 \pm 2.7$) | 66 ($\pm 18.1$) |
| **Sex (female**<br>**%)** | 7 (36.8%) | 29 (36%) | 12 (38%) | 17 (34%) |
| **Weight (kg)** | 75.5 ($\pm 18.1$) | 76.4 ($\pm 17.7$) | 76.3 ($\pm 15.9$) | 79.8 ($\pm 17.6$) |
| **Height (cm)** | 173.3 ($\pm 8.3$) | 173.2 ($\pm 9.4$) | 173.5 ($\pm 11.1$) | 173.3 ($\pm 7.0$) |
| **BMI (kg/m<sup>2</sup>)** | 28.2 ($\pm 3.6$) | 26.3 ($\pm 4.8$) | 25.9 ($\pm 5.5$) | 26.3 ($\pm 3.3$) |
| **AFLD** | nA | 42 | 16 | 50 |
| **NAFLD** | nA | 26 | 9 | nA |
| **Hepatitis B** | nA | 5 | 1 | nA |
| **Hepatitis C** | nA | 4 | 3 | nA |
| **PBC/PSC/AIH** | nA | 2 | 2 | nA |
| **MELD** | nA | 10.8 ($\pm 5.7$) | 12.7 ($\pm 3.3$) | 19.2 ($\pm 5.5$) |
| **Leucocytes**<br>**(g/l)** | 7.2 ($\pm 1.6$) | 5.0 ($\pm 2.3$) | 5.9 ($\pm 2.5$) | 5.8 ($\pm 2.8$) |
| **Hemoglobin**<br>**(g/l)** | 140.5 ($\pm 16.3$) | 123.0 ($\pm 22.6$) | 122.2 ($\pm 21.3$) | 109.0 ($\pm 20.4$) |
| **Hematocrit**<br>**(%)** | 0.4 ($\pm 0.05$) | 0.35 ($\pm 0.06$) | 0.35 ($\pm 0.05$) | 0.31 ($\pm 0.06$) |
| **Platelet count**<br>**(g/l)** | 272.9 ($\pm 66.4$) | 128.8 ($\pm 80.4$) | 100.3 ($\pm 51.5$) | 79.5 ($\pm 61.8$) |
| **Urea (mg/dl)** | 34.2 ($\pm 16.3$) | 33.4 ($\pm 17.8$) | 43.9 ($\pm 24.1$) | 59.4 ($\pm 31.3$) |
| **Creatinine**<br>**(mg/dl)** | 0.9 ($\pm 0.1$) | 0.8 ($\pm 0.4$) | 1.2 ($\pm 0.9$) | 1.2 ($\pm 0.8$) |
| **Filtration rate**<br>**(ml/min/m<sup>2</sup>)** | 58.2 ($\pm 9.2$) | 57.0 ($\pm 10.3$) | 55.4 ($\pm 8.4$) | 56.4 ($\pm 9.1$) |
| **Bilirubin**<br>**(mg/dl)** | 0.8 ($\pm 0.4$) | 1.4 ($\pm 0.7$) | 4.2 ($\pm 2.6$) | 5.7 ($\pm 2.3$) |
| **Sodium**<br>**(mmol/l)** | 140.4 ($\pm 2.7$) | 138 ($\pm 4.2$) | 137.0 ($\pm 5.8$) | 135.1 ($\pm 5.8$) |
| **AST (U/l)** | 36.6 ($\pm 13.2$) | 60.7 ($\pm 31.2$) | 88.5 ($\pm 39.9$) | 98.2 ($\pm 41.4$) |
| **ALT (U/l)** | 29.1 ($\pm 17.2$) | 44.9 ($\pm 28.0$) | 71.0 ($\pm 58.2$) | 89.1 ($\pm 28.5$) |
| **Gamma-GT**<br>**(U/l)** | 63.5 ($\pm 34$) | 101.0 ($\pm 59.5$) | 182.3 ($\pm 52.8$) | 222.5 ($\pm 55.6$) |
| **Alkaline Phosphatase (U/l)** | 73.6 (±33.8) | 143.0 (±47.8) | 220.2 (±150.5) | 145.0 (±85.0) |
| **LDH (U/l)** | 179.0 (±43.8) | 244.3 (±99.0) | 260.0 (±132.1) | 240.1 (±63.8) |
| **CRP (mg/dl)** | 0.5 (±0.2) | 1.1 (±0.5) | 1.5 (±0.8) | 1.8 (±0.8) |
| **Quick (%)** | 99.0 (±13.3) | 67.3 (±17.3) | 55.9 (±22.6) | 54.5 (±15.8) |
| **INR** | 1.0 (±0.1) | 1.3 (±0.2) | 1.4 (±0.5) | 1.5 (±0.4) |
| **PT (sec)** | 30.5 (±2.6) | 38.7 (±9.1) | 46.1 (±17.3) | 43.2 (±18.4) |
| **Fibrinogen (mg/dl)** | 334.0 (±53.5) | 274.4 (±106.8) | 212.9 (±144.5) | 204.6 (±83.1) |
| **Albumin (mg/dl)** | 4170.7 (±399.9) | 3362.7 (±607.4) | 3090.8 (±591.4) | 3014.8 (±476.2) |
| **Alpha-1-Fetoprotein (µg/l)** | 1.8 (±0.4) | 7.4 (±4.2) | 6.9 (±3.8) | 7.4 (±3.4) |

Data are expressed as case numbers (percentage) or mean ± standard deviation. kg – kilograms, cm – centimeter, kg/m<sup>2</sup> – kilograms/square meter, AFLD – alcoholic fatty liver disease, NAFLD – nonalcoholic fatty liver disease, PBC – primary biliary cholangitis, PSC – primary sclerosing cholangitis, AIH – autoimmune hepatitis, mg/dl – milligrams/deciliter, G/l – grams/liter, mmol/l – millimole/liter, ml/min/m<sup>2</sup> – milliliters/minute/square meter, U/l – Units/liter, sec – second, µg/l – microgram/liter

Further exclusion criteria were uncontrolled infection, multiorgan failure, uncontrolled upper gastrointestinal bleeding, preexisting kidney disease with serum creatinine > 2.5 mg/dl, hepatocellular carcinoma (HCC) or other active malignancies, pregnant and lactating people and active drug abuse. In the AH group patients with hepatitis B or C, acquired human immune deficiency syndrome, autoimmune liver disease, primary biliary cholangitis and primary sclerosing cholangitis, Wilson disease, hemochromatosis, and suspected drug-induced liver injury were excluded.

Healthy volunteers, with no underlying disease, (n=19) were included in our outpatient clinic during regularly check-up (n=15). 4 healthy volunteers underwent liver biopsy as potential living liver donors, no liver disease was detected histologically.

Patients included in the validation cohort (n= 186) distributed as followed: 23 healthy volunteers, 85 patients suffering from compensated cirrhosis, 42 patients suffering from decompensated cirrhosis and 36 patients with AH contributed to the validation patient cohort. Further details of the patient cohort are presented in Table 2.

**Table 2:**
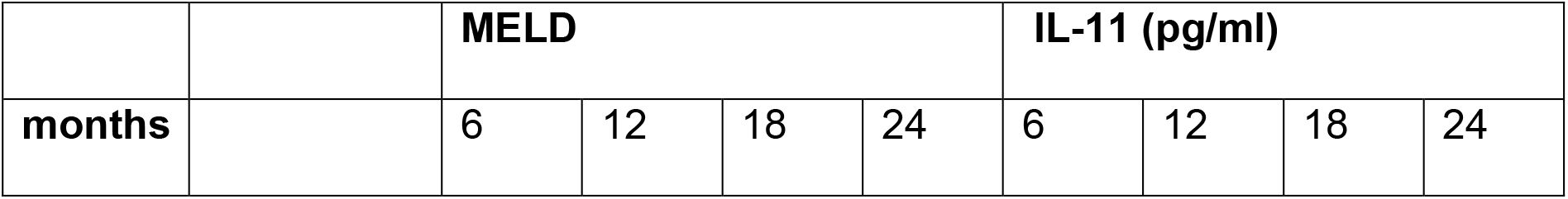
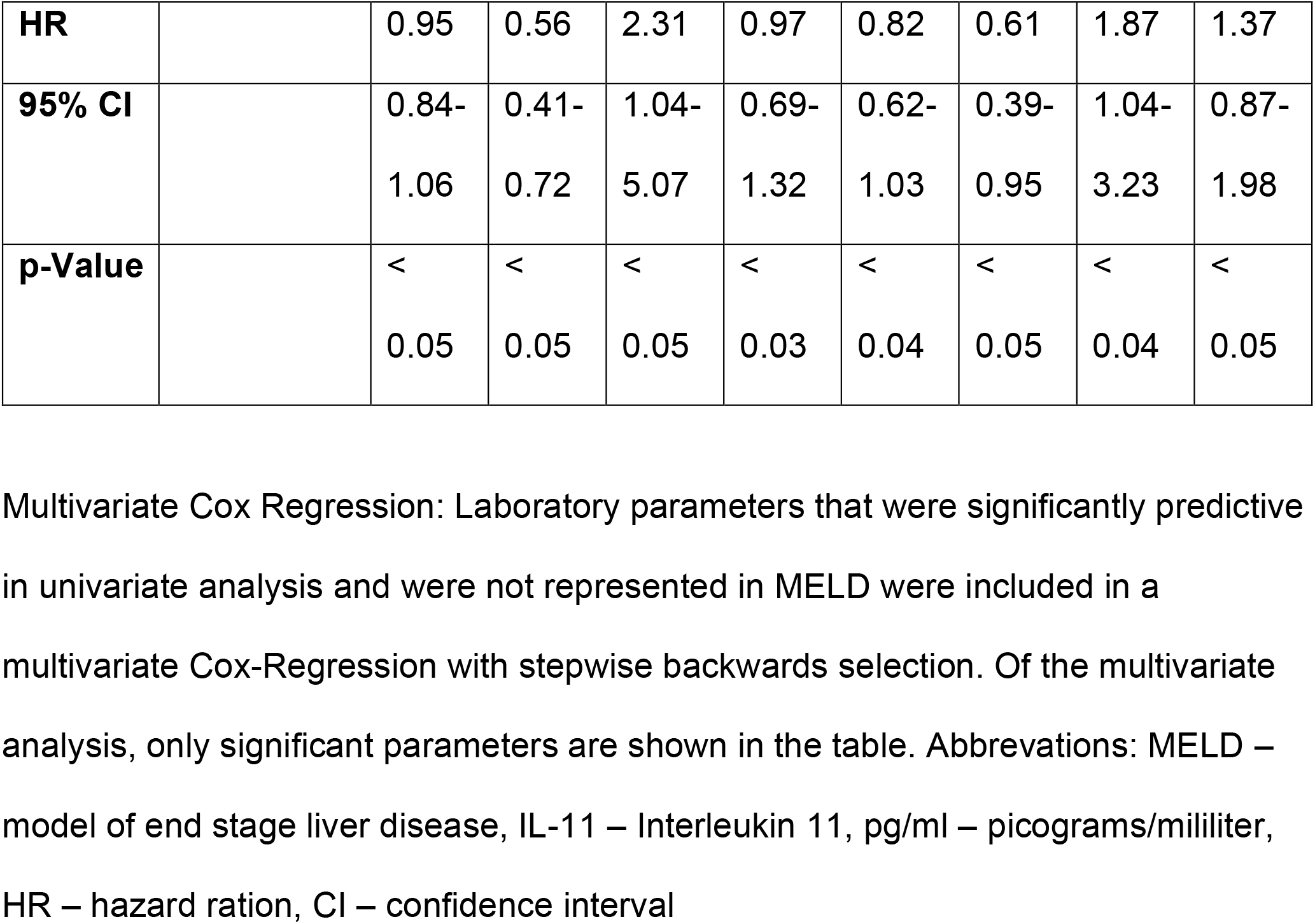
Multivariate COX Regression.

### Ethical consideration

The study protocol was approved by the institutional ethics commission with an amendment to AN2017-0016 369/4.21.

### Mouse studies

C57BL/6 mice purchased from Charles River Laboratories (Wilmington, MA) were cohoused in the Animal Facility of Medical University of Innsbruck for one week prior to the start of experiments. All mice were fed the Lieber DeCarli pair-fed diet for five days to become acclimated to a liquid diet. Female wild-type (wt) mice (7-8 weeks old) were then fed with a Lieber-DeCarli diet (BioServ, Flemington, NJ) containing an increasing amount of ethanol (EtOH) ranging from 1 to 5 vol% ad libitum for 15 days (EtOH-fed)^22^. Control diet was supplemented with an isocaloric amount of maltose (pair-fed). Pair-fed mice were calorie matched with the ethanol-fed mice. Mice were weighed every other day. 8 hours after gavage the mice were euthanized. All mice received Xylain 5 mg/kg bodyweight (Intervet, Vienna, Austria) and Ketamin 100 mg/kg bodyweight (AniMedica, Senden, Germany) for anesthesia. Blood, tissue samples of liver and intestine were collected afterwards. Serum was stored at −80°C as well as tissue samples were stored at −80°C or in RNAlater (Qiagen, Hilden, Germany) at −20°C.

### In Vivo administration of anti-IL11RA

Mice were injected intraperitoneally with either anti-IL11RA (kindly provided by the Cook laboratory, Duke-NUS, Singapore) or an identical amount of Immunoglobulin G (IgG) control (20 mg/kg) (kindly provided by the Cook laboratory, Duke-NUS, Singapore). These antibodies have been described in detail in previous publications^18,23^

### Ethical considerations

All anti-IL11RA experiments adhered to ethical principles according to Austrian law (BMWFW-2020.0.547.764) and the ARRIVE Guidelines^24^. The experiments were carried out at the animal facility of the Medical University of Innsbruck.

### Data analysis

Data are expressed as mean ± standard error of mean or as median with first and third quartiles. For comparing quantitative variables, the Student’s t-test or the non-parametric Mann–Whitney U or Wilcoxon signed-rank test were used as appropriate.

Normality of distribution was determined by Kolmogorov-Smirnov test. The correlation analysis was estimated using the Spearman’s p coefficient. Survival analysis was performed using the Kaplan-Meier method applying the log-rank test for group comparison. Kruskal-Wallis test or ANOVA was used where applicable. To identify predictors of mortality, a Cox proportional hazards model was applied. Correlation was carried out using the Spearman’s rank correlation coefficient; - to correct for multiple testing, the Benjamin-Hochberg method was used with an accepted false discovery rate of < 0.05. A p-value < 0.05 was considered as statistically significant. All statistical analyses were performed using SPSS Statistics v.22 (IBM, Chicago, IL), R Programming and GraphPad PRISM 5 (La Jolla, CA).

Further information on materials and methods are provided in the supplementary material and methods section.

## Results

Our training cohort included 160 patients with liver disease being treated in-hospital or in our outpatient clinic, with etiologies listed in Table 1. The cohort comprised 79 patients with compensated cirrhosis and 31 patients with decompensated cirrhosis of varying etiology, while 50 patients participated with AH. The study design is depicted in Fig. 1A. To exclude any preexisting liver diseases in patients suffering from alcoholic hepatitis different staining methods have been used, e.g. Masson’s trichrome staining or periodic acid shift with diastase staining (Supplementary Fig. 1). Healthy volunteers (n=19) served as control group, with characteristics provided in Table 1.

**Fig. 1:**
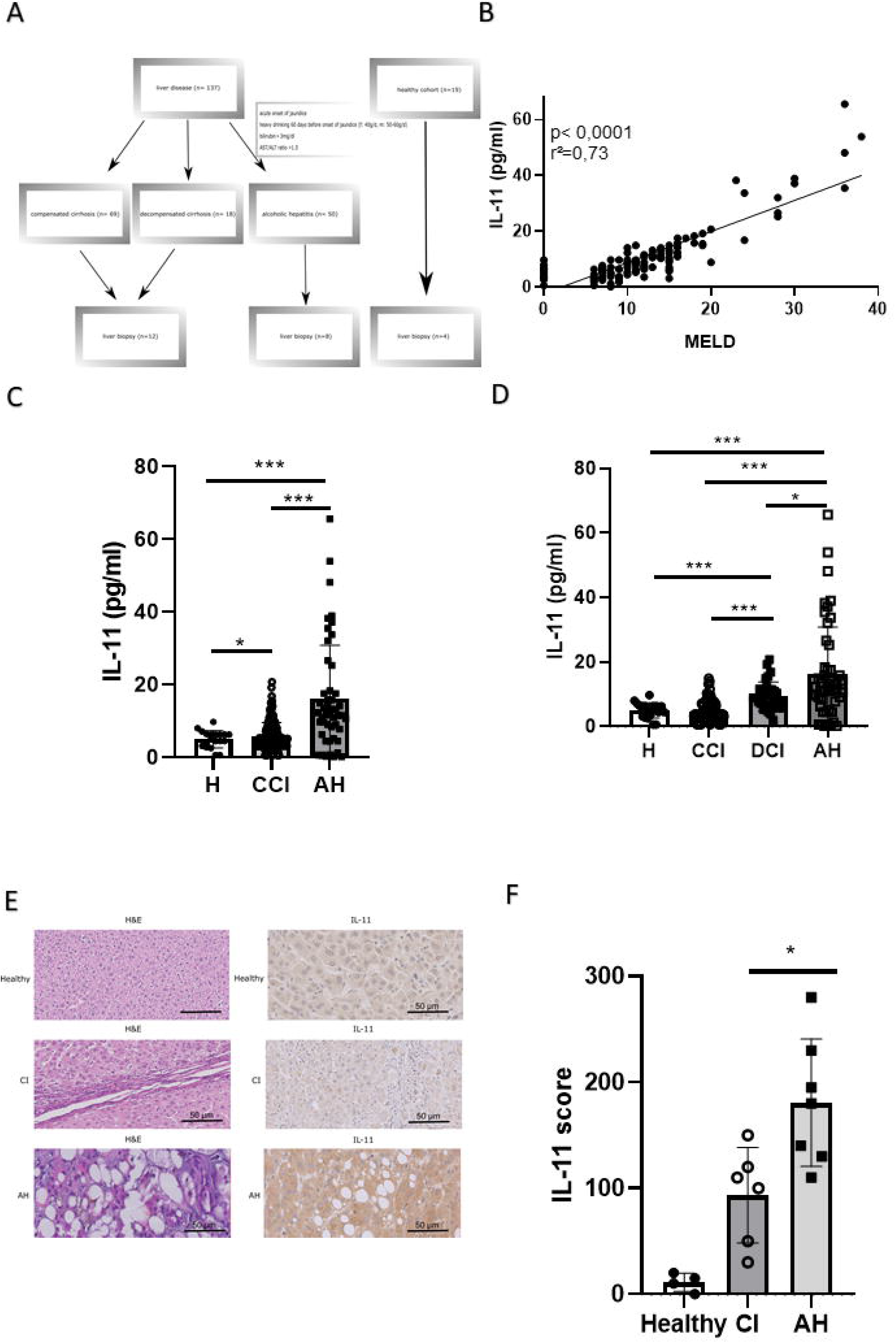
Severity of ALD correlates with circulating IL-11 levels and its hepatic expression. Study design and patient’s selection (A). IL-11 serum levels and MELD score correlation in patients suffering from cirrhosis and alcoholic hepatitis and healthy controls (MELD in healthy controls was assumed 0) (B) IL-11 levels correlate with cirrhosis and alcoholic hepatitis in patients compared to healthy controls (MELD in healthy controls was assumed 0) (C). IL-11 levels distinguish between patients with compensated or decompensated cirrhosis, alcoholic hepatitis and healthy controls. (MELD in healthy controls was assumed 0) (D). Representative HE staining and IL-11 staining in liver tissue in healthy controls, patients suffering from cirrhosis and patients suffering from alcoholic hepatitis (E). Statistical analysis of IL-11 positive cells in liver tissue (F). (\**p* <0.05, \*\**p* <0.01, \*\*\**p* <0.001) Abbreviations: IL-11 – Interleukin 11, MELD – model of end stage liver disease, pg/ml – picograms/milliliters, H - healthy control, CI – cirrhosis, CCI – compensated cirrhosis, DCI – decompensated cirrhosis, AH – alcoholic hepatitis, µm – micrometer, HE – hematoxylin and eosinophils.

### Severity of ALD correlates with circulating IL-11 levels and its hepatic expression

The MELD score positively correlated with IL-11 levels (p<0.001, r^2^=0.73, Fig. 1B) and IL-11 was significantly elevated in cirrhosis and AH patients compared to healthy controls, respectively. Highest levels were seen in alcoholic hepatitis (Fig. 1C). IL-11 serum levels were higher in decompensated cirrhosis compared to compensated patients (Fig. 1D). There was no correlation of IL-11 levels with C-reactive protein, albumin, INR, liver transaminases, gamma-glutamyl transferase, alkaline phosphatase, leukocytes, and serum creatinine (Supplementary Fig. 2). Interestingly, etiology of liver cirrhosis, excluding AH, was not reflected in different IL-11 levels (data not shown).

To examine IL-11 levels at the level of liver tissue, hematoxylin and eosin (HE) and immunohistochemistry (IHC) IL-11 staining was performed in liver biopsies from healthy volunteers, patients with cirrhosis and alcoholic hepatitis (Fig. 1E). A large number of IL-11 positive hepatocytes were detected in ongoing AH, whereas biopsies of cirrhotic liver tissue stained less for IL-11 expression (Fig. 1F).

### Serum IL-11 concentration predicts patient outcome

Univariate and multivariate Cox regression analyses were performed to uncover non-invasive prediction markers for survival up to 24 months without liver transplantation. After Cox regression analyses, non-significant factors like age or gender and factors already represented in the MELD analysis were excluded based on stepwise model selection. Finally, MELD and IL-11 concentration remained in our multivariate model and both turned out to independently predict survival without transplantation for 6, 12, 18, and 24 months (Table 2). The accuracy of IL-11 levels in predicting transplant-free survival was comparable to the established MELD score in a receiver operating characteristics (ROC) analysis (Fig. 2A). Youden’s index was calculated to determine an optimal cut-off value for IL-11 for liver disease. IL-11 below 6.4 pg/ml predicted a transplant-free 6 months survival with a sensitivity of 59.4% and a specificity of 93.3% (Fig. 2A). To further evaluate the performance of IL-11 levels, we analyzed the role of IL-11 serum levels in a training cohort (n=179) using the Kaplan-Meier method (p-value of 0.0001) (Fig. 2B). Data were checked and compared in a validation cohort (n=186) (Fig. 2C). Characteristics of the validation cohort are summarized in Supplementary Table 2. Notably, transplant-free survival was significantly lower in the validation cohort (validation cohort vs. training cohort 28%; p<0.001). Similar as in the training cohort, the accuracy of IL-11 to predict transplant free survival was non - inferior to that of MELD (Supplementary Figure 3). Furthermore IL-11 AUROCs in the training and validation cohort didn’t show significant differences (Supplementary Figure 3). Differences between the AUROCs in the training and in the validation cohort were not statistically significant (Supplementary Table 3).

**Fig. 2:**
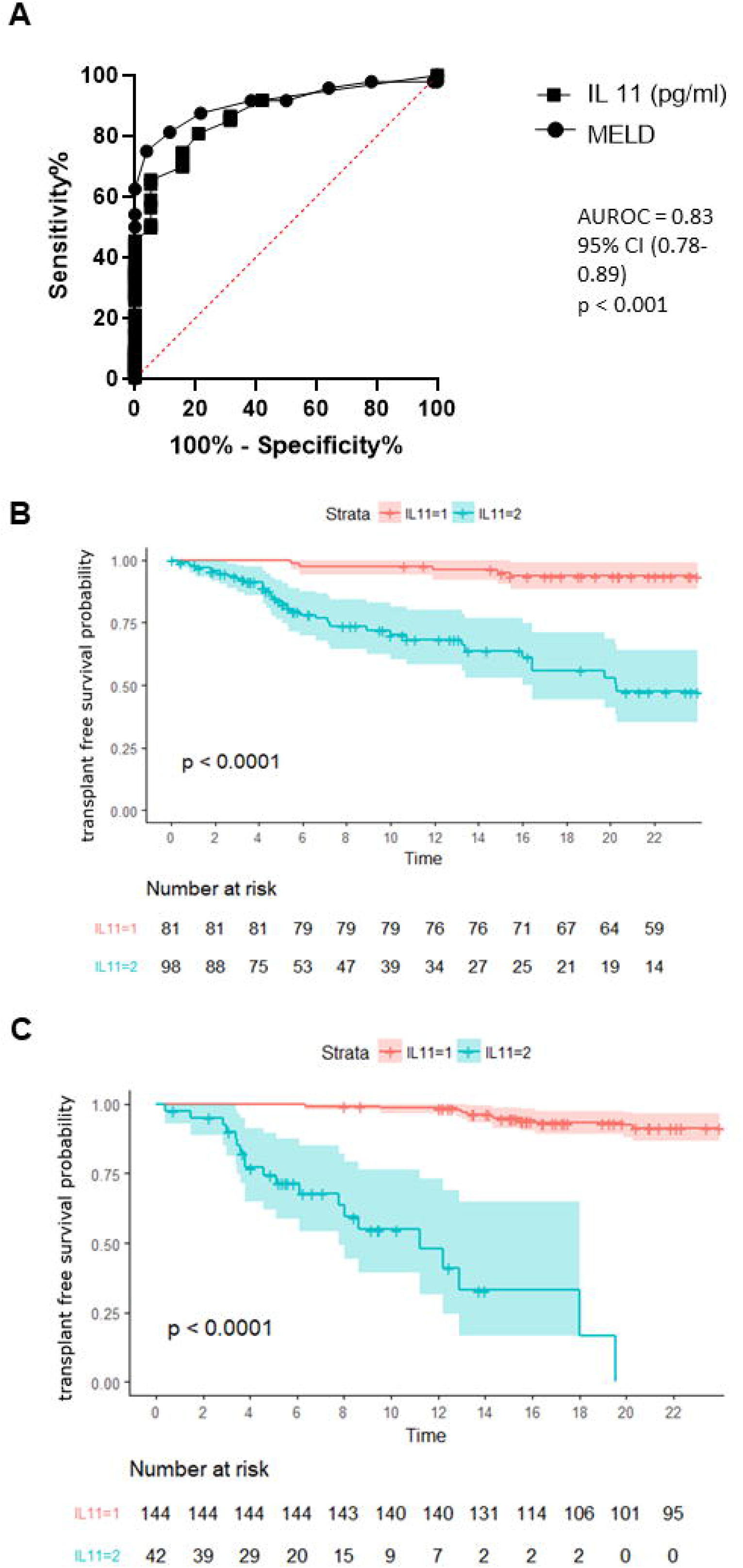
Serum IL-11 concentration predicts patient outcome. AUROCs of the scores for prediction of transplant free survival (A). Kaplan Meyer curve for transplant-free survival with IL-11 cutoff 6.4 pg/ml (B). Kaplan Meier curve for validation cohort (C). IL-11=1: Group 1 with serum IL-11 concentration < 6.4 pg/ml. IL-11=2: Group 2 with serum IL-11 concentration > 6.4 pg/ml. Abbreviations in order of their appearance: ALD – alcoholic liver disease, IL-11 – Interleukin 11, MELD – model of end stage liver disease, pg/ml – picograms/milliliters.

### Inhibition of IL-11 signaling protects against experimental ALD

Female C57BL/6J mice were exposed to a 5% ethanol containing Lieber-DeCarli diet or an isocaloric pair diet for 15 days. In this experimental set-up EtOH-fed mice received a neutralizing IL-11 receptor antibody (anti-IL11RA) or IgG control intraperitoneally as illustrated in Fig. 3A. Control-treated EtOH-fed mice showed signs of liver injury which was reversed by anti-IL11RA administration (Fig. 3B). Furthermore, anti-IL11RA administration resulted in reduced liver-body ratio compared to IgG control (Fig. 3C) and anti-IL11RA could also prevent the EtOH-induced weight loss (Fig. 3D). The anti-IL11RA effect applied also to EtOH-induced hepatic triglyceride accumulation (Fig. 3E).

**Fig. 3:**
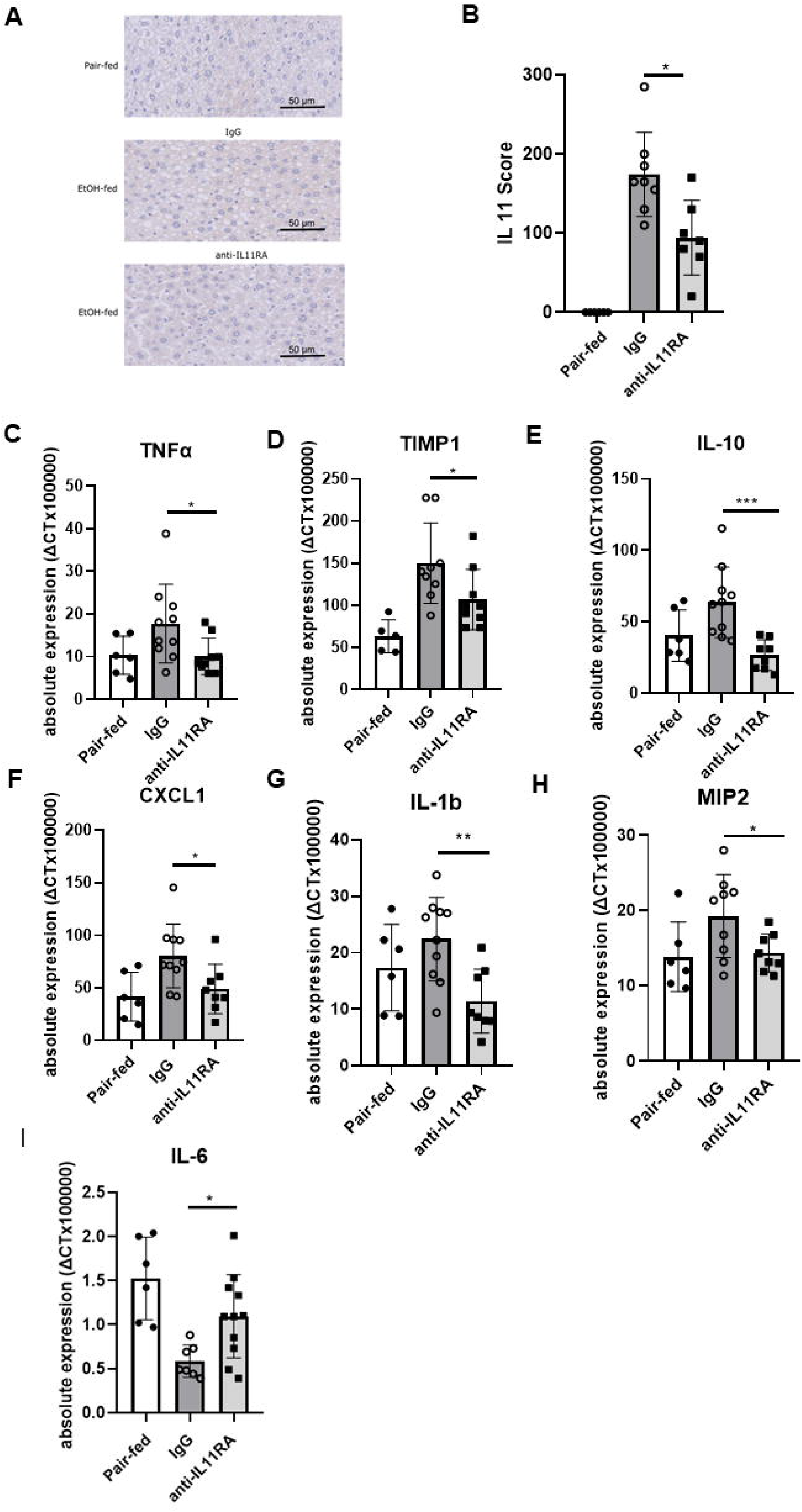
Inhibition of IL-11 signaling protects against experimental ALD. Schematic timeline of the procedure. C57BL/6J female mice were pair-fed or EtOH-fed for 15 days. The EtOH-group received either IgG or anti-IL11RA intraperitoneally (A). Application of anti-IL11RA resulted in significant lower ALT levels in the EtOH group compared to the IgG group (B). EtOH fed mice treated with anti-IL11RA showed decreased liver-to-body ratio (C), decreased weight loss (D) and decreased hepatic triglyceride accumulation compared to EtOH fed mice treated with IgG (E). (\**p* <0.05, \*\**p* <0.01, \*\*\**p* <0.001, n ≥5/group) Abbreviations in order of their appearance: anti-IL11RA – Interleukin 11 Receptor Antibody, EtOH – Ethanol, IgG – Immunoglobulin G, ALT – alanine transferase, IL-11 – Interleukin 11

### Inhibition of IL-11 signaling protects against ethanol-induced liver inflammation

To investigate any further potential beneficial effects of the anti-IL11RA, we stained hepatic tissue of these mice for IL-11 (Fig. 4A) showing significant lower expression of IL-11 in anti-IL11RA-treated compared to ethanol-fed control mice (Fig. 4B). Since IL-11 might induce other pro-inflammatory mediators in the liver we further investigated hepatic gene expression of TNFα, tissue inhibitor of metalloproteinases 1 (TIMP-1), IL-10, CXCL1, IL-1ß, macrophage inflammatory proteins (MIP)-2 and IL-6. Anti-IL11RA treatment significantly reduced expression of each of them apart from IL-6, which has been reported to be potentially beneficial in liver disease ^25^ (Figure 4C – 4I). Remarkably, the expression of IL-1β was suppressed below baseline. Additional inflammation and fibrosis marker are shown in Supplementary Fig. 4.

**Fig. 4:**
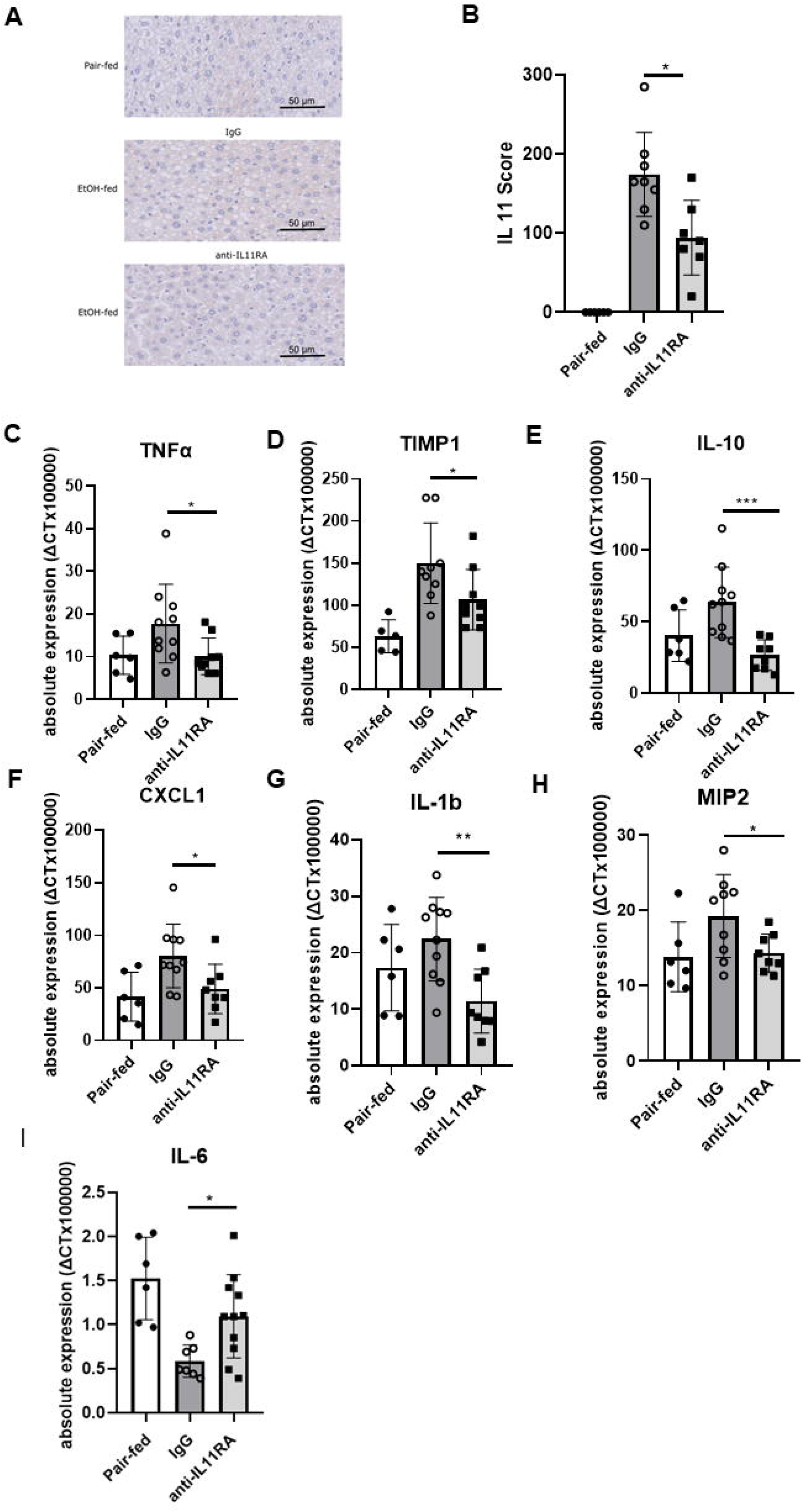
Inhibition of IL-11 signaling protects against ethanol-induced liver inflammation. Representative IL-11 staining images from all 3 groups (A). Administration of IL-11RA tend to reduce IL-11 in liver tissue (B). Effect of IL-11RA administration on hepatic expression of proinflammatory cytokines in EtOH fed mice: TNF-α (C), TIMP1 (D), IL-10 (E), CXCL-1 (F), IL-1b (G), MIP 2 (H) and IL-6 (I). (\**p* <0.05, \*\**p* <0.01, \*\*\**p* <0.001, n ≥5/group) Abbreviations in order of their appearance: IL-11R– Interleukin-11 receptor, IL-11 – Interleukin 11, anti-IL11RA – Interleukin 11 Receptor Antibody, EtOH – Ethanol, IgG – Immunoglobulin G, TIMP1-metallopeptidase inhibitor 1, IL-10 – Interleukin-10, TNFα – tumor necrosis factor – α, CXCL-1 – chemokine (C-X-C motif) ligand 1, IL-1β – interleukin 1 beta, MIP 2 – macrophage inflammatory protein 2

### Anti-IL11RA treatment reduces infiltration of pro-inflammatory cells into the liver

After HE-staining of liver tissue 20 HPFs were analyzed, and a steatosis score was calculated. Steatosis score was significantly reduced upon treatment with anti-IL11RA compared to control animals (Fig. 5AB). Myeloperoxidase (MPO+) staining showed an almost complete inhibition of neutrophil immigration into liver tissue by systemic anti-IL11RA treatment (Fig. 5CD). This was consistent with decreased F4/80 positive macrophages in ethanol-fed mice treated with anti-IL11RA (Fig. 5EF).

**Fig. 5:**
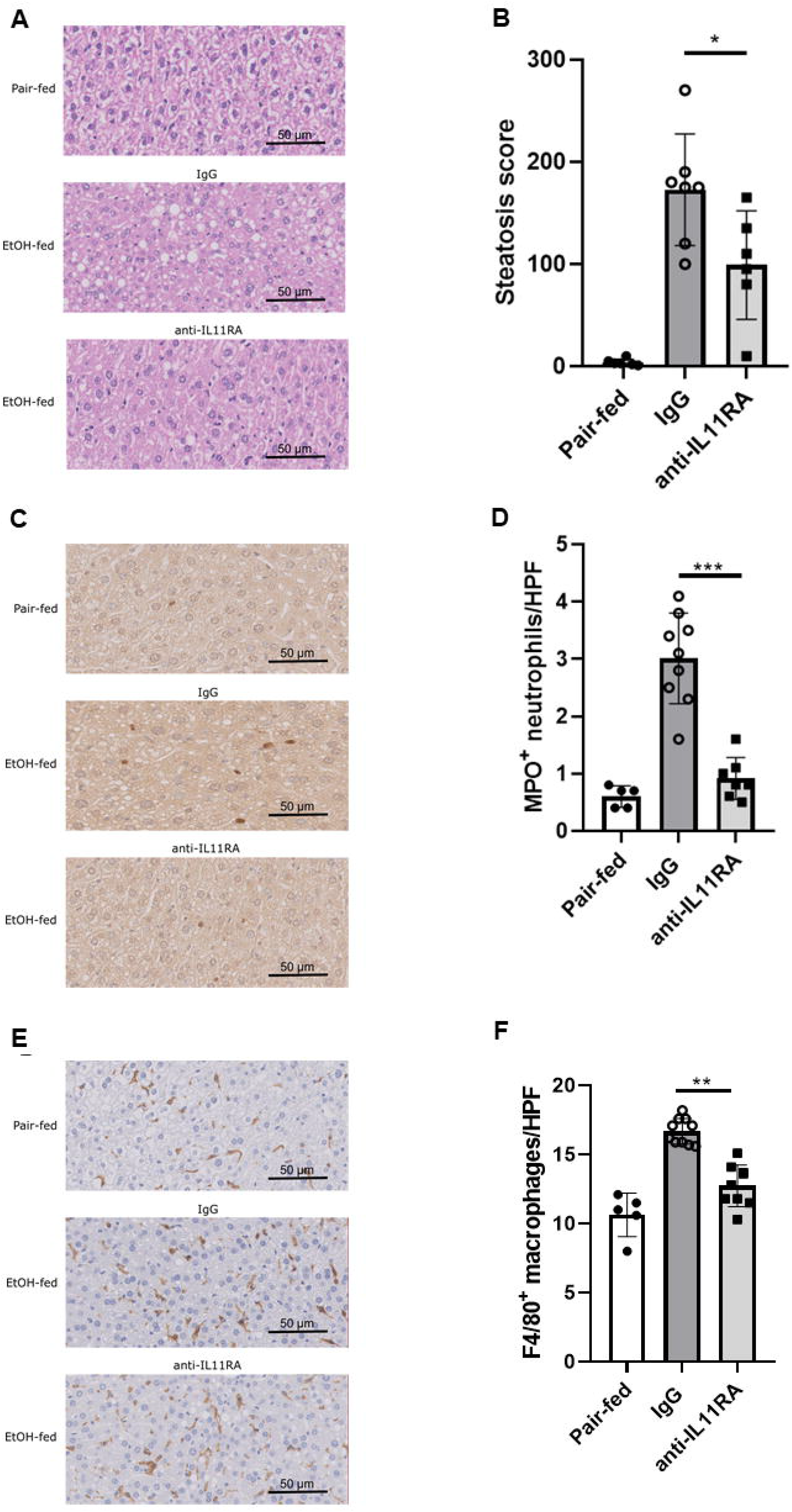
Anti-IL11RA treatment reduces infiltration of pro-inflammatory cells into the liver. Representative H&E staining images from all 3 groups (A). Hepatic steatosis score decreased significantly in EtOH-fed wt mice treated with IL-11RA (B). Representative images and quantification of MPO^+^ cells per high power field in the liver based on hepatic MPO immunoreactivity (D). Hepatic infiltration of MPO^+^ neutrophils are significantly decreased in EtOH-fed mice treated with anti-IL11RA (E). Representative images and quantification of F4/80 positive macrophages per high power field in the liver based on hepatic F4/80 immunoreactivity (D). Hepatic infiltration of F4/80 macrophages are significantly decreased in EtOH-fed mice treated with anti-IL11RA (F). n ≥5/group. Abbrevations in order of their appearance: anti-IL11RA – Interleukin 11 Receptor Antibody, IgG – Immunoglobulin G, EtOH – Ethanol, HPF – high power field, H&E – hematoxylin and eosin, MPO^+^ - myeloperoxidase positive

## Discussion

We herein report increased IL-11 levels in peripheral blood along with upregulated intrahepatic IL-11 expression in patients suffering from advanced liver diseases, specifically in patients with AH. A positive correlation was found with transplant-free survival and a cut off value for IL-11 serum levels was determined and validated in a separate cohort. We assumed IL-11 might play an important role in the hepatotoxicity of ethanol and/or pro-inflammatory processes and that blocking IL-11 receptor may have beneficial effects on multiple components of ALD. This hypothesis is supported by our murine ALD model using a neutralizing IL-11 receptor antibody.

Parenchymal infiltration of neutrophils and macrophages is a prominent feature of ALD and is likely due to ethanol-mediated activation of innate immunity and subsequent induction of pro-inflammatory cytokines and chemokines^26-29^. Recent studies discovered an unexpected pro-inflammatory role for IL-11 in NAFLD and showed that HSCs highly express IL-11 receptor and show an indirect effect of IL-11 on immune cells that is mediated via the stroma^18^. It is also the case that injured hepatocytes, which also highly express IL11RA, secrete IL-11 and this activates HSCs and inflammatory cells in a paracrine manner while damaging hepatocytes in an autocrine fashion ^13^. We could corroborate these findings in our study in the context of ALD. In line with this, anti-IL11RA robustly reduced inflammation in our ALD model. Hepatocytes express the IL-11 receptor and secrete cytokines upon ligation, such as transforming growth factor beta (TGF-β)^18^. IL-11 activation of hepatocytes is unexpectedly cytotoxic and an autocrine and maladaptive loop of IL-11 activity in hepatocytes is apparent. The effects of IL-11 on these cell type during acute necroinflammation in non-alcoholic steatohepatitis (NASH) are profound^13^ and were also observed in our ALD model. IL-11 blockade was also associated with lower liver fat and reduced hepatic oxidative stress in murine NASH models^30^.

However, former studies have suggested that IL-11 might act cytoprotective, anti-fibrotic, and anti-inflammatory based on effects of recombinant human IL-11 in mouse models of hepatic disease^31-35^. These findings led to a clinical trial using rhIL-11 in patients suffering from hepatitis C but this study was not taken forward^36^. The suggested protective role of IL-11 was questioned by recent animal studies showing that species-matched IL-11 is a key player in liver fibrosis in NAFLD as well as in the development of severe inflammation in NASH^13,18,37^.

Widjaja and co-workers recently showed that species-matched IL-11 is hepatotoxic and induces reactive oxygen species (ROS)-dependent hepatocyte cell death via c-Jun N-terminal kinase (JNK) along with inhibition of liver regeneration^18^. The difference of these findings with the former literature, where a high dose of rhIL-11 was injected to rodents, may be explained by the fact that rhIL-11 binds to the mouse IL-11 receptor, but it does not activate the same signaling pathways as endogenous murine IL-11 does. Dong et al. recently showed that IL-11 modulates the hepatocytic metabolism and suggested a transition mechanism via the cis-pathway from NAFLD to NASH, conditions where IL-11 levels are elevated in the liver and serum ^13^. In *Ilra1* knockout mice, restoration of IL-11 cis-signaling in hepatocytes reestablished steatosis and inflammation^13^. The pro-inflammatory effect could be confirmed in our murine model of ALD, but most importantly, was also seen in patients with AH. Compared to cirrhosis patients, the circulating IL-11 levels were much higher. These data strongly suggest an important role of the IL-11 pathway in pathogenesis of human AH, which is consistent with the more recent literature on IL-11.

In a mouse model of APAP induced liver damage i.e., Widjaja et al. showed that circulating IL-11 levels are markedly elevated and genetic or pharmacologic inhibition of IL-11 signaling reduced liver damage and promoted liver regeneration^23^. These findings are in line with elevated IL-11 concentrations as a pro-inflammatory marker in ALD and NASH. Notably, in NASH, genetic or antibody-induced inhibition of IL-11 signaling reduced inflammation ^13^. These findings are also in accordance with our findings that support further the idea that IL-11 as a pro-inflammatory cytokine in murine ALD.

Our study has some limitations. While the National Institute on Alcohol Abuse and Alcoholism (NIAAA) mouse model provides excellent insight into liver inflammation, liver fibrosis cannot be sufficiently addressed in this model^22^. We found a beneficial effect of IL-11 inhibition on liver steatosis and neutrophilic inflammation. We could consistently show effects of IL-11 inhibition on pro-inflammatory factors, but we did not specifically investigate effects on immune cells themselves or the effects of endotoxins on liver derived cytokines, chemokines and reactive oxygen species (ROS). In addition, our training and validation cohort were collected from the same region/hospital.

In conclusion, IL-11 plays a key role in inflammation and liver damage in our murine model of ALD and we showed elevated IL-11 levels in blood and liver tissue in human AH and liver cirrhosis of various etiologies, with a clear cut-off to predict outcome. We propose that inhibition of IL-11 signalling as a key factor for reducing inflammation and hepatotoxicity in alcoholic liver disease might also impact long term fibrosis development. Therefore, anti-IL-11RA therapy could be a promising therapeutic option in treating AH and preventing cirrhosis.

## Abbrevations

AH: alcoholic hepatitis
ALD: alcoholic liver disease
ALT: alanine aminotransferase
anti-IL11RA: Interleukin 11 receptor antagonist
AST: aspartate transaminase
ETOH: ethanol
HE: hematoxylin and eosin
HCC: hepatocellular carcinoma
HSC: hepatic stellate cells
IgG: Immunglobulin G
IHC: immunohistochemistry
IL: Interleukin
JNK: c-Jun N-terminal kinase
MELD: Model of End stage Liver Disease
MIP: macrophage inflammatory protein
NAFLD: non-alcoholic fatty liver disease
NASH: non-alcoholic steatohepatitis
NIAAA: National Institute on Alcohol Abuse and Alcoholism
RA: receptor antibody
ROC: receiver operating characteristic
ROS: reactive oxygen species
TNF-α: tumor necrosis factor-α
TGFβ: transforming growth factor beta
wt: wildtype

